# Correlation between the immuno-virological response and the nutritional profile of treatment-experienced HIV-infected patients in the East region of Cameroon

**DOI:** 10.1101/2020.02.11.943621

**Authors:** Abba Aissatou, Joseph Fokam, Rachel Simo Kamgaing, Junie Flore Yimga, Aude Christelle Ka’e, Alex Durand Nka, Michel Carlos Tommo Tchouaket, Ambe Collins Chenwi, Ezechiel Ngoufack Jagni Semengue, Alexis Ndjolo, Samuel Martin Sosso

## Abstract

**Background:** HIV management remains concerning and even more challenging in the frame of comorbidities like malnutrition that favors disease progression and mortality in resource-limited settings (RLS).

**Objective:** To evaluate the correlation between immuno-virological responses and the nutritional profile of HIV-infected individuals receiving antiretroviral therapy (ART).

**Methods:** A cross-sectional study was conducted from October to December 2018 among 146 consenting participants enrolled in two health facilities of the East-Region of Cameroon. Socio-demographic data, basic clinical information and treatment history were collected; blood samples were collected by venipuncture for laboratory analysis (HIV-1 viral load, CD4-CD8 Tcells measurement and biochemical analysis) performed at the “Chantal Biya” International Reference Center”, Yaounde, Cameroon. Nutritional profile was evaluated using anthropometric and biochemical parameters. Data were analyzed using Excel 2016, Graph pad prism version 6 and R.version3.5.0; Spearman correlation was used; with p<0.05 considered statistically significant.

**Results:** Median [IQR] age was 42 [33-51] years, 76.0% (111/146) were female and median [IQR] duration on ART was 54 [28-86] months. Of these participants, 11.6% (17/146) were underweight based on the body mass index and 4.7% (7/146) were at the stage of advanced weight loss. According to immunovirological responses, 44.5% (65/146) were immunocompromised (CD4<500 cell/µl) and 75.3% (110/146) had an undetectable viremia (<40 copies/mL). CD4 count inversely correlated with total protein concentration (r=-0.18, p=0.030) and viremia was inversely correlated with total cholesterol (r=-0.65; p=0.001), and positively correlated with total protein (r=0.28; p<0.001) and seemingly with triglycerides (r=0.27; p=0.070) concentrations.

**Conclusion:** In this RLS with patients having about five years of ART-experience, half are immunocompromised while the majority have achieved good virological response. Interestingly, one out of eight patients might be experiencing malnutrition. Specifically, increasing CD4 may favour hypo-proteinemia while increasing viral load may prone hyper-proteinemia and hypo-cholesterolemia. Further studies are needed in RLS with high burden of HIV-infection.

## Introduction

The human immunodeficiency virus (HIV) targets the immune system and weakens the surveillance and defense system of the body against infections, leading to susceptibility of HIV infected individuals to a wide range of infections normally cleared by the immune system of a healthy/immunocompetent individual [1]. HIV can therefore cause several health complications including opportunistic infections, oxidative stress, wasting syndrome, as well as malnutrition [2].

Malnutrition is one of the major complications of HIV infection [3] and has been recognized under the banner of ‘wasting syndrome’ as a significant prognostic factor of advanced disease [4]. Of note, malnutrition is defined as unhealthy diets deficient in micronutrients and micronutrient imbalances, which can disrupt the function of various immune system [5]. Specifically, under-nutrition impairs the immune system mechanism and thus impairs the host response against micro-organisms. The consequence of this impairment is an increase in both incidence and severity of infections [6]. Particularly, HIV infection and insufficient nutritional intake are part of a vicious cycle that contributes to immunodeficiency and poor health outcomes [7].

HIV/AIDS and malnutrition have a synergistic interaction within the host. In effect, malnutrition increases the risk of HIV pathogenesis, while HIV in turn triggers malnutrition by depleting the immune system from nutrient intake, absorption and utilization [8]. There is a complex triangulation mechanism between malnutrition, the immune system and HIV infection, in which malnutrition elicits immune system dysfunctions which in turn promotes increased vulnerability of the host to infection, while the latter intensifies the severity of malnutrition [9].

Progress in scaling-up HIV treatment (23.3 million, representing 62% global coverage) have reduced associated mortality and morbidity, thus making HIV a chronic infection, even in sub-Saharan African countries where 70% of the global epidemics is concentrated [10–13]. Thus, for an enhanced performance of ART programs, it is postulated that nutritional interventions should become an integral component in the management of people living with HIV (PLHIV). Such strategy suggests that improved attention to diet and nutrition could normalize protein profiling, fatty acids, copper and iron, which in turn would harness the immune function and subsequently optimize ART acceptability, adherence and effectiveness [14,15].

In spite of the declining burden of HIV at country-level (5.4% in 2004 to 2.7% in 2019), Cameroon is still experiencing a generalized epidemiology [16–18]. The national strategy for the fight against HIV/AIDS in Cameroon recommends a nutritional supplement to subside the current 14.1% persistent rate of malnutrition for PLHIV, and in turn to support ART response [19,20]. Interestingly, the East region of Cameroon has a very high burden of HIV infection (5.6%), 43.01% patients are on ART [18,21], while up to 13.7% of these patients are known to be undernourished [20]. Thus, these high rates of both HIV infection and malnutrition call for further investigations to improve the management of this comorbidity in similar settings of sub-Saharan Africa.

We therefore sought to study the correlation between the immuno-virological response and the nutritional profile among ART-experienced HIV-infected patients in the East-region of Cameroon.

## Materials and methods

### Study design, sampling method and eligibility criteria

A cross-sectional and analytical study was carried-out among PLHIV receiving ART at two heath facilities in the East region of Cameroon, Bertoua Regional Hospital (BRH); and Nkolbikon Catholic Health Center (NCHC), during three months (from October through December 2018).

Following a consecutive sampling, the required minimum sample size for the study was calculated based on the prevalence of undernourished PLHIV in the East-region of Cameroon (13.7%) [20], as per the following statistical formula: 

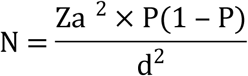

With “Z” equal to 1.96 at 95% confidence interval, with “P” the prevalence of undernourished PLHIV in our study setting (13.7%) [20], and “d” being the error rate set at 6% (0.06); after numerical application, we obtained a minimum sample size, “N” = 126.2, rounded-up to a minimum of 127 participants to be enrolled in the study.

Eligibility criteria were every individual: 1) with confirmed HIV-positive; 2) on ART for at least six (06) months; 3) registered in one of the study sites; and 4) aged 15 years and above. Following these criteria, a total of 146 participants were enrolled.

### Phlebotomy and sample shipment

A standard questionnaire was administered to each participant by interviewers trained on the study protocol. Blood sample was then collected (in dry and Ethylenediaminetetraacetic acid test tubes of 4 ml) by venipuncture with the help of trained phlebotomists. After collection, blood samples were transferred from the sampling sites to the BRH laboratory for packaging and shipment. Only dry test tubes were centrifuged and separated in aliquots for biochemical analysis. After centrifugation, racks of all samples were placed in ice packed isothermal bags, and samples were then sent for laboratory analysis to the study reference institution (Chantal BIYA International Reference Center for Research on HIV/AIDS prevention and management), located in Yaoundé, the capital city of Cameroon.

### Clinical and laboratory procedures

During routine clinic attendance at the study sites, the standard questionnaire (S1 file) administered to all participants enabled us to retrieve information covering socio-demographic data, treatment history and food habits, as well as basic clinical information and all biological parameters performed.

CD4 and CD8 cell count were performed using the Cyflow Counter-Sysmex Partec as per the manufacturer’s instructions with results reported as the number of positive cells per microliter of blood (http://www.nsplucknow.com/pdf/CyFlow_Counter_Bro_EN.pdf). CD4 results were then interpreted as follows: no immunodeficiency (above or equal to 500); moderate immunodeficiency (between 350 and 499); advanced immunodeficiency (between 200 and 349) and severe immunodeficiency (below 200) [22]. Regarding the interpretation of CD8 cell results: no immunodeficiency was defined as absolute value above 220 cells while immunodeficiency was below 220, as previously described [23]. Viral load measurement was performed using the Abbott m2000RT Real Time PCR as-per the manufacturer’s instructions (Abbott Laboratories, USA), with a lower detection threshold of 40 HIV-1 RNA copies/mL and an upper detection threshold of 10,000,000 copies/mL (www.abbottmolecular.com/products/infectious-diseases/realtime-pcr/hiv-1-assay).

The nutritional profile of participants was evaluated based on biochemical parameters: albumin, calcium, glucose, iron, magnesium, total cholesterol, triglycerides, total protein and anthropometric parameters of which the body mass index (BMI), nutritional risk index (NRI) and weight loss percentage (WLP). Biochemical analysis was performed using BT-3000 Plus as per manufacturer’s instructions (https://www.chema.com/chema/automation_it_files/Biotecnica%20BT3000.pdf). BMI was defined as an indicator of chronic energy malnutrition and was calculated by dividing the weight (in Kg) by the height squared (in square meter): 

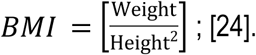

The NRI, a nutritional status assessment index recommended in national nutritional programs, was estimated by the following formula: 

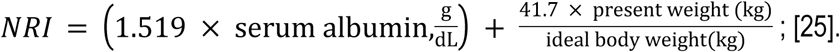

WLP is used to assess the risk of malnutrition and determined by the formula 

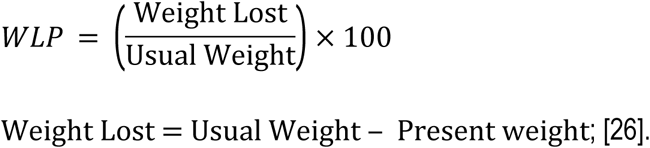

### Statistical Analysis

Collected data were entered on Microsoft Excel 2016 sheets and analyzed using the software GraphPad Prism version 6.0. Dependent variables were immune status and viremia, while independent variables were anthropometric and biochemical parameters. Correlation analysis were performed using Spearman correlation test; with p<0.05 considered statistically significant. The software R version 3.5.0. was used for backward multiple regression with AIC (Akaike Information Criteria) selection. P-values ≤0.2, obtained in bivariate analysis, were considered for the multivariate analysis.

### Ethical Considerations

The study was conducted in compliance with the core principles of the Helsinki declaration: an administrative authorization was issued; ethical clearance was obtained from the National Ethics Committee for Research on Human Health (ref N^0^ 2018/06/1055/CE/CNERSH/SP); written informed consent were obtained from all the participants; data were processed using unique identifiers to ensure confidentiality; laboratory results were returned to participants for possible benefits in their clinical management; and counseling on good nutritional habits and healthy lifestyles was provided to all.

## Results

### Socio-demographic and clinical characteristics of the population

Of the total of 146 participants enrolled, 88.4% (129) at Bertoua regional hospital (BRH) and 11.6% (17) at Nkolbikon Catholic Health Center (NCHC). Among them, 76.0%(111) were females giving a sex ratio (female/male) of 3. The median age was 42 [IQR: 33-51] years (Table 1).

The median duration on ART was 54 [IQR: 28-86] months. All participants were on first-line ART regimen consisting of two nucleoside reverse transcriptase inhibitors (NRTIs) and one non-NRTI (NNRTI).

**Table 1:** Socio-demographic data

| Variables | Effective | Percentage (%) |
| --- | --- | --- |
| **Health Center** |  |  |
| BRH | 129 | 88.4 |
| NCHC | 17 | 11.6 |
| **Gender** |  |  |
| Male | 35 | 23.9 |
| Female | 111 | 76.0 |
| **Economic status per day (Francs CFA)** |  |  |
| <500 | 8 | 5.5 |
| 500-1000 | 27 | 18.5 |
| 1000-5000 | 107 | 73.3 |
| >5000 | 4 | 3.7 |
| **Average of meals per day** |  |  |
| 1 meal | 12 | 8.2 |
| 2 meals | 72 | 49.3 |
| 3 meals | 54 | 36.9 |
| 4 meals | 8 | 5.5 |

**Legend:** BRH: Bertoua regional hospital; NCHC: Nkolbikon Catholic Health Center

### Nutritional profile

Of the 146 participants, BMI results revealed that only a proportion of 11.6% (17) was undernourished. According to NRI, 91.1% (133) were normal, while referring to WLP, 60.9% (89) had a normal weight. Only 2.7% (4) of participants suffered from hypoalbuminemia, 19.9% (29) from hyper-proteinemia and 20.5%(30) from hypo-cholesterolemia (Table 2).

**Table 2:**
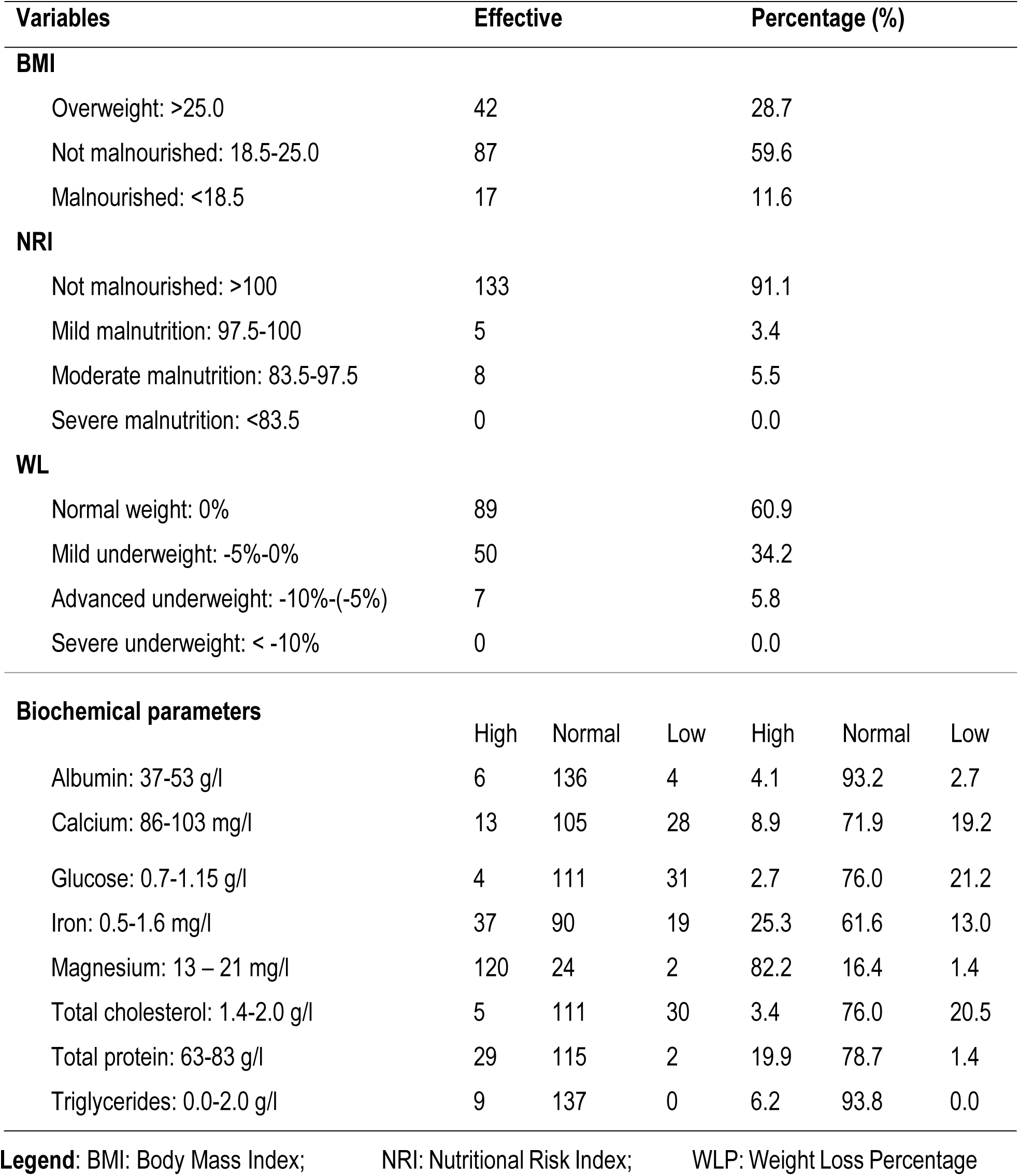
Anthropometric and biochemical parameters

| Variables | Effective |  |  | Percentage (%) |  |  |
| --- | --- | --- | --- | --- | --- | --- |
| **BMI** |  |  |  |  |  |  |
| Overweight: >25.0 | 42 |  |  | 28.7 |  |  |
| Not malnourished: 18.5-25.0 | 87 |  |  | 59.6 |  |  |
| Malnourished: <18.5 | 17 |  |  | 11.6 |  |  |
| **NRI** |  |  |  |  |  |  |
| Not malnourished: >100 | 133 |  |  | 91.1 |  |  |
| Mild malnutrition: 97.5-100 | 5 |  |  | 3.4 |  |  |
| Moderate malnutrition: 83.5-97.5 | 8 |  |  | 5.5 |  |  |
| Severe malnutrition: <83.5 | 0 |  |  | 0.0 |  |  |
| **WL** |  |  |  |  |  |  |
| Normal weight: 0% | 89 |  |  | 60.9 |  |  |
| Mild underweight: -5%-0% | 50 |  |  | 34.2 |  |  |
| Advanced underweight: -10%-(-5%) | 7 |  |  | 5.8 |  |  |
| Severe underweight: < -10% | 0 |  |  | 0.0 |  |  |
| **Biochemical parameters** |  |  |  |  |  |  |
|  | High | Normal | Low | High | Normal | Low |
| Albumin: 37-53 g/l | 6 | 136 | 4 | 4.1 | 93.2 | 2.7 |
| Calcium: 86-103 mg/l | 13 | 105 | 28 | 8.9 | 71.9 | 19.2 |
| Glucose: 0.7-1.15 g/l | 4 | 111 | 31 | 2.7 | 76.0 | 21.2 |
| Iron: 0.5-1.6 mg/l | 37 | 90 | 19 | 25.3 | 61.6 | 13.0 |
| Magnesium: 13 – 21 mg/l | 120 | 24 | 2 | 82.2 | 16.4 | 1.4 |
| Total cholesterol: 1.4-2.0 g/l | 5 | 111 | 30 | 3.4 | 76.0 | 20.5 |
| Total protein: 63-83 g/l | 29 | 115 | 2 | 19.9 | 78.7 | 1.4 |
| Triglycerides: 0.0-2.0 g/l | 9 | 137 | 0 | 6.2 | 93.8 | 0.0 |

**Legend:** BMI: Body Mass Index; NRI: Nutritional Risk Index; WLP: Weight Loss Percentage

### Immuno-virological status

The median [IQR] CD4+ T cell lymphocyte count was 547 [385-759] cells/μl, and 44.5% (65) were immunocompromised (<500 cells/μl). The median [IQR] CD8+ T lymphocytes was 692 [512-997] cells/μl, and only one had a value below 220 cells cells/μl. According to CD4+/CD8+ ratio, 26.0% (38) of our population were immunocompromised (ratio<0.5). The majority (75.3%) of the 146 study participants had a controlled of viral replication (<40 copies/mL) while 12.3% (18) were experiencing virological failure (>1000 copies/mL) (Table 3).

**Table 3:** Immuno-virological results

|  | N | % |
| --- | --- | --- |
| CD4 cell (cells/μl) |  |  |
| Below 200 | 4 | 2.7 |
| Between 200-349 | 25 | 17.1 |
| Between 350-499 | 36 | 24.6 |
| Above 500 | 81 | 55.5 |
| CD8 cell (cells/μl) |  |  |
| Below 220 | 1 | 0.7 |
| Above 220 | 138 | 99.3 |
| CD4/CD8 |  |  |
| Below 0.5 | 38 | 26.0 |
| Between 0.5-2.0 | 106 | 72.6 |
| Between 2.0 | 2 | 1.4 |
| ART received |  |  |
| 3TC+TDF+EFV | 123 | 84.2 |
| 3TC+TDF+EFV+Cotrim | 2 | 1.4 |
| 3TC+AZT+NVP | 15 | 10.3 |
| 3TC+ABC+EFV | 2 | 1.4 |
| 3TC+AZT+EFV | 4 | 2.7 |
| Viremia (copies/mL) |  |  |
| Viral control (undetectable) | 110 | 75.3 |
| Low level viremia (40-1000) | 18 | 12.3 |
| Virological failure (>1000) | 18 | 12.3 |

**Legend:** 3TC: Lamivudine; ABC: Abacavir; ART: Antiretroviral Therapy; AZT: Zidovudine; CD: Cluster of Differentiation; EFV: Efavirenz; NVP: Nevirapine; TDF: Tenofovir

### Correlation between immuno-virological parameters and nutritional profile

We found a negatively weak correlation between CD4+ T cells count and total protein concentration (r =-0.18; p=0.03) as shown on figure 1. CD8+ showed a negatively weak correlation with glucose concentration (r=-0.24; p=0.003) and a positively weak correlation with total protein concentration (r=0.18; p=0.031) as shown in figure 2 and figure 3 respectively. Bivariate analysis of viremia with each nutritional parameter are presented in table 4. Viral load showed a negatively weak correlation between with total cholesterol (r=-0.28, p=0.0007), and a positively weak correlation with total protein (r= 0.28, p=0.0006) and with triglycerides (r= 0.20, P=0.018), as shown figure 4, figure 5 and figure 6 respectively. After multivariate analysis including BMI, WLP, NRI, total cholesterol, total protein, triglycerides and albumin, three parameters (total cholesterol, total protein and triglycerides) were found to be independently correlated to viral load, as shown on table 5.

**Figure 1:**
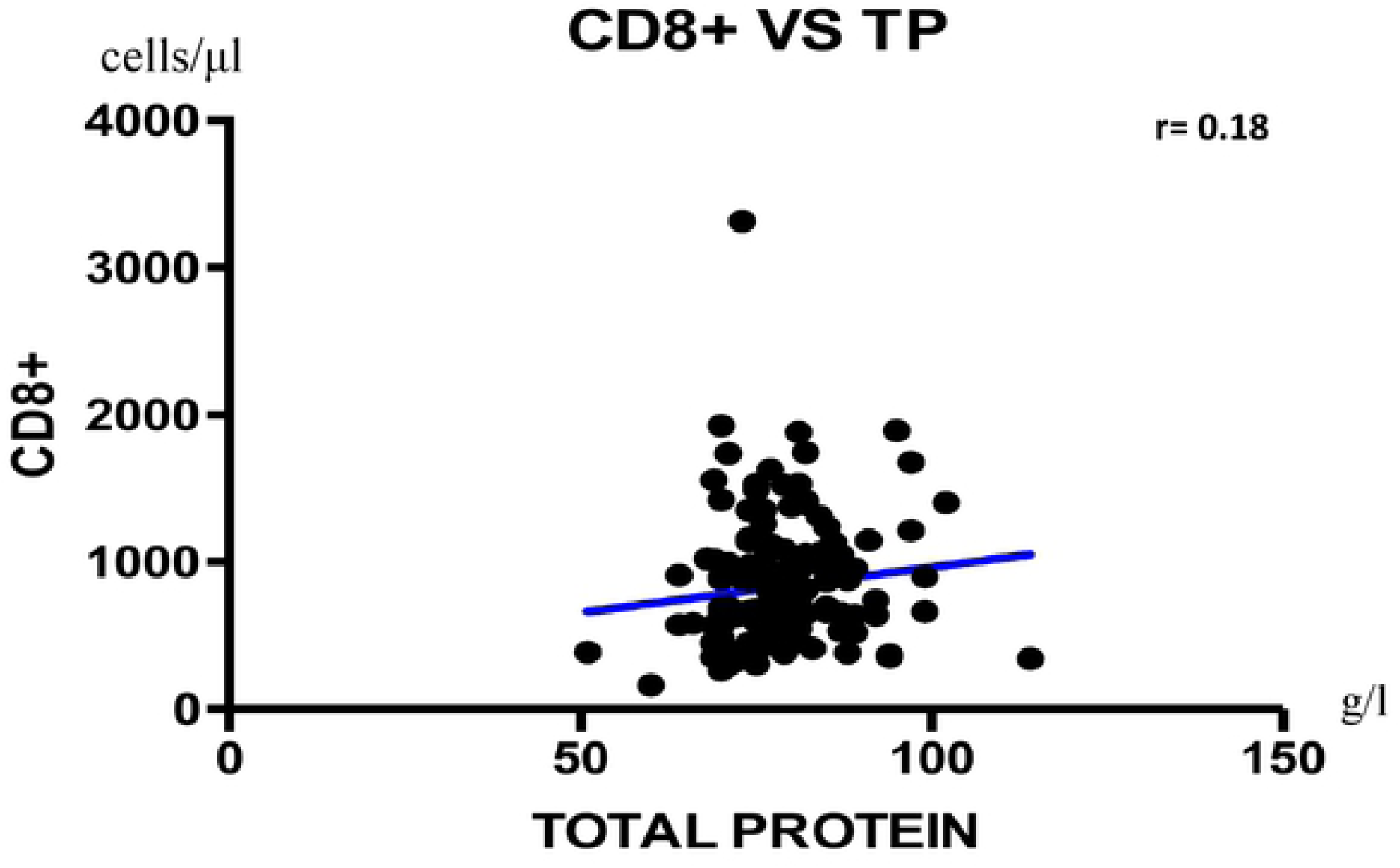
Correlation between LTCD4+ and Total protein.

**Figure 2:**
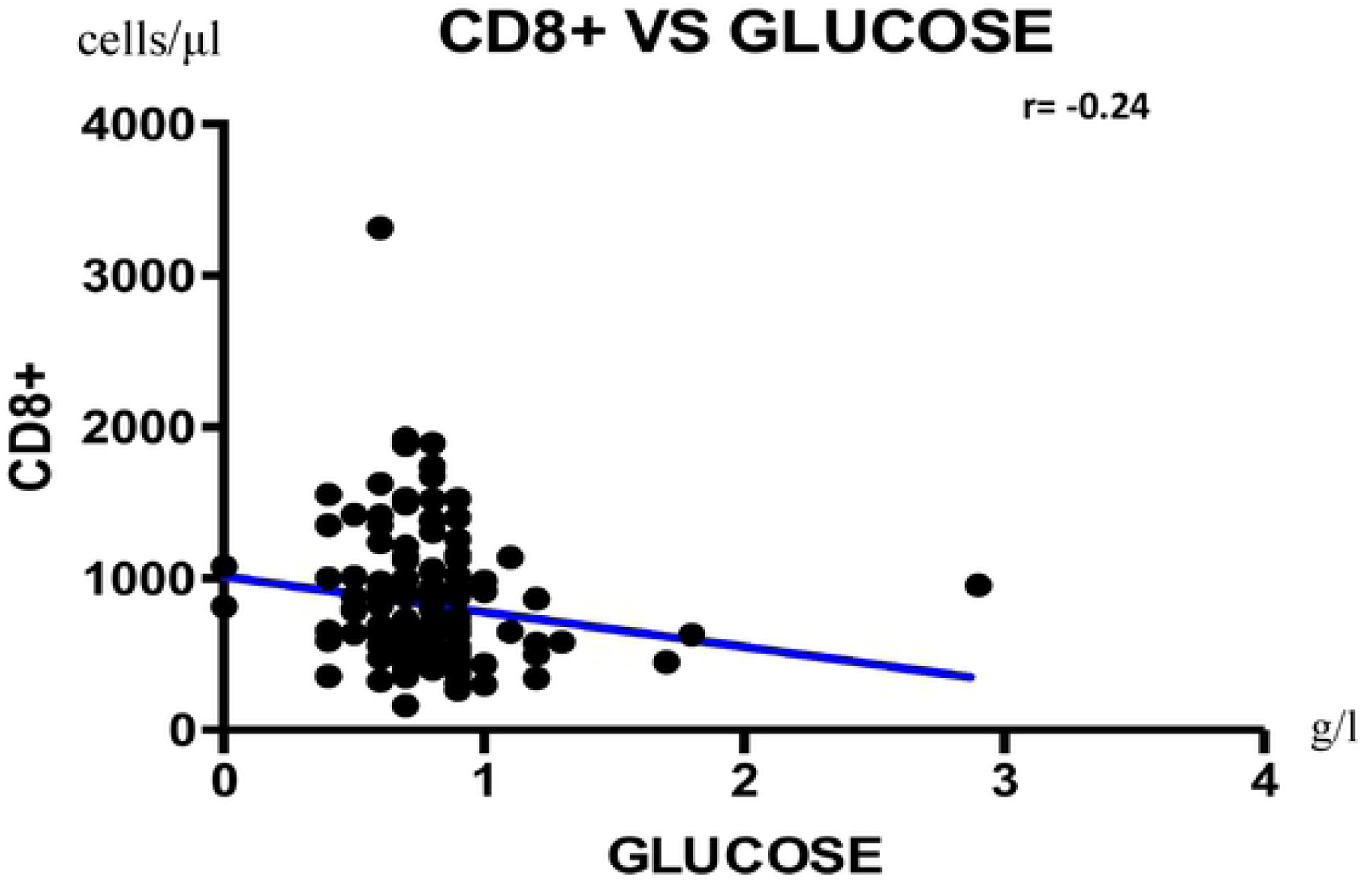
Correlation between LTCD8+ and Total Protein.

**Figure 3:**
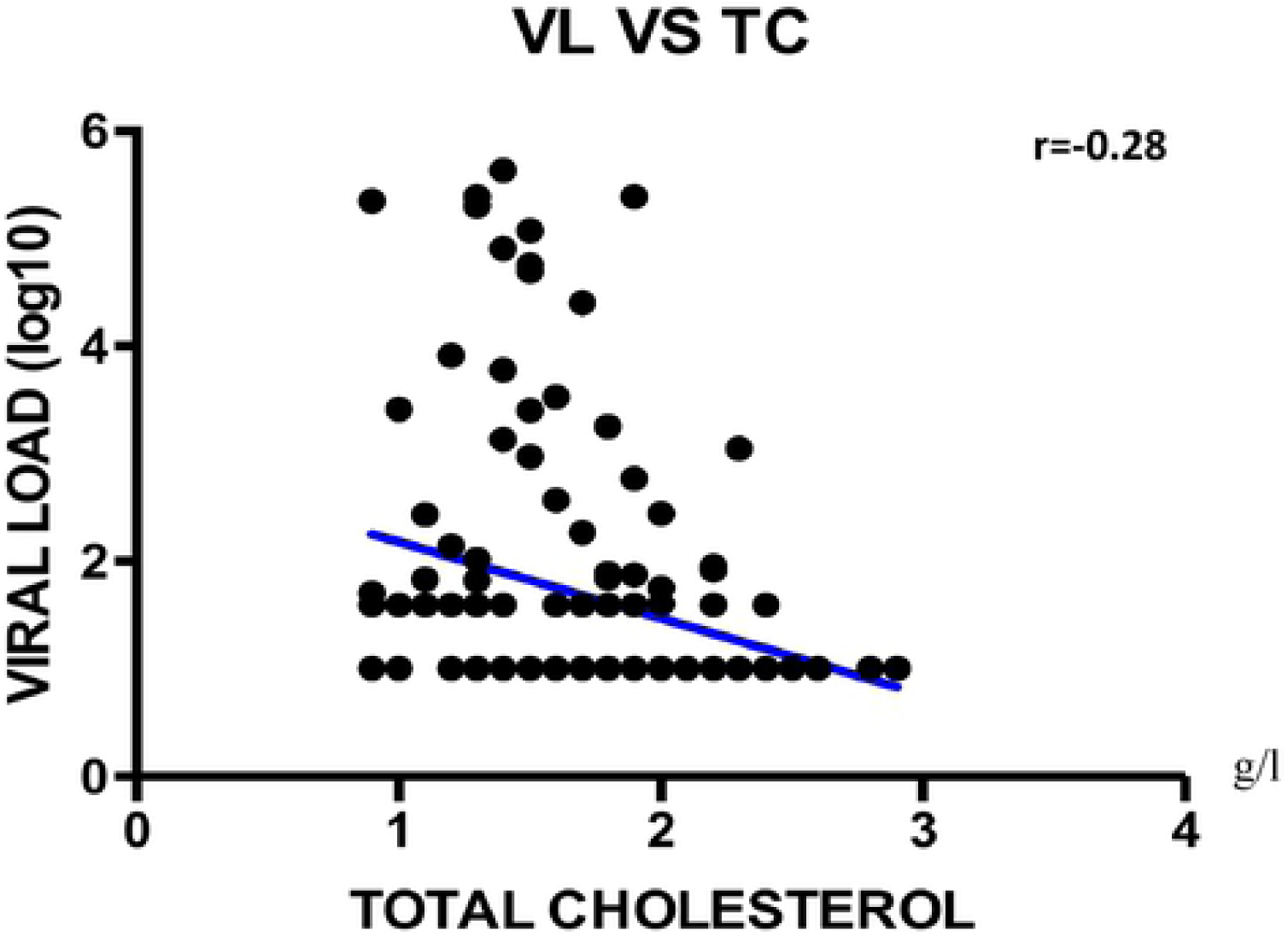
Correlation between LTCD8+ and Glucose.

**Figure 4:**
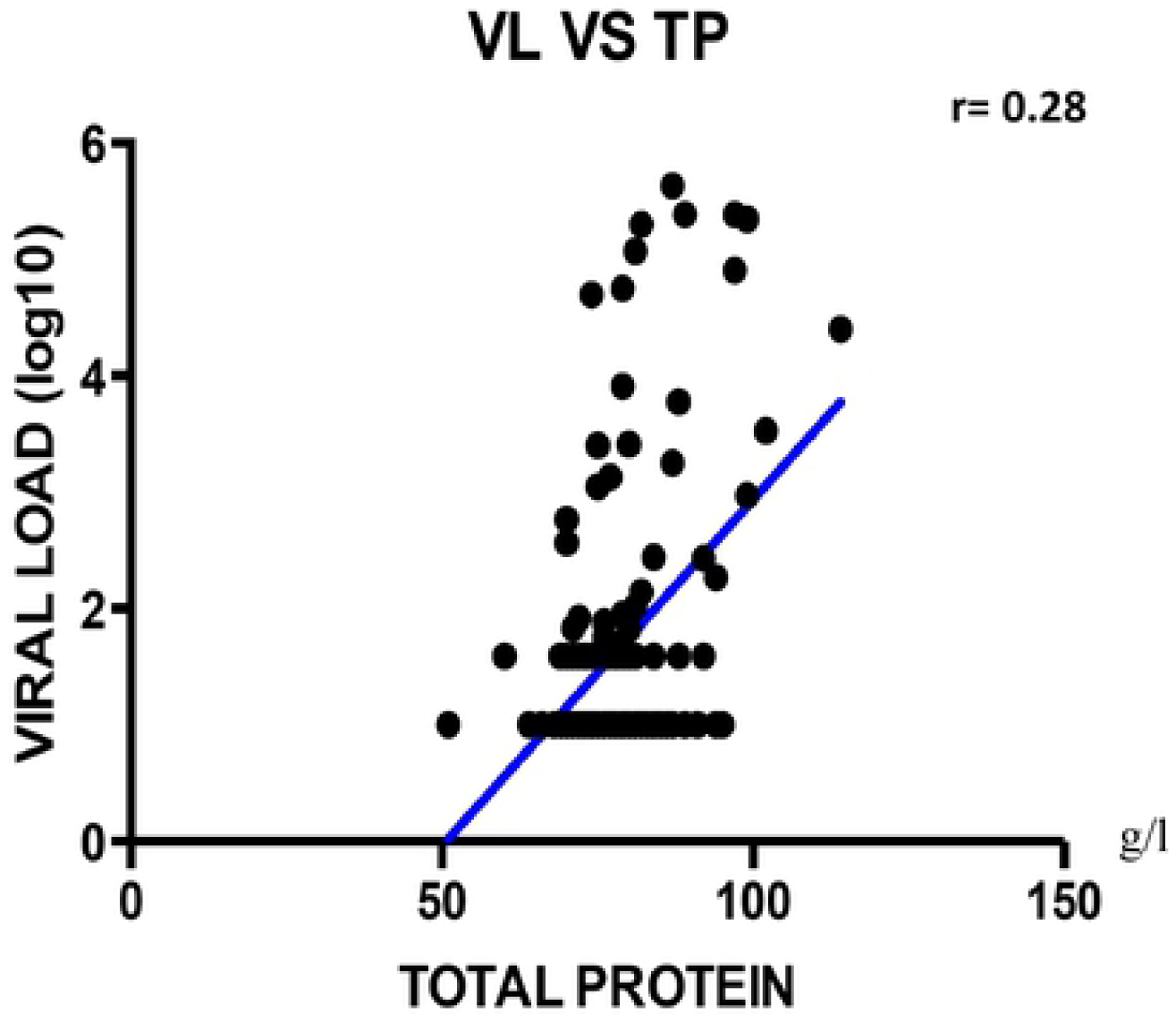
Correlation between viral load and total cholesterol.

**Figure 5:**
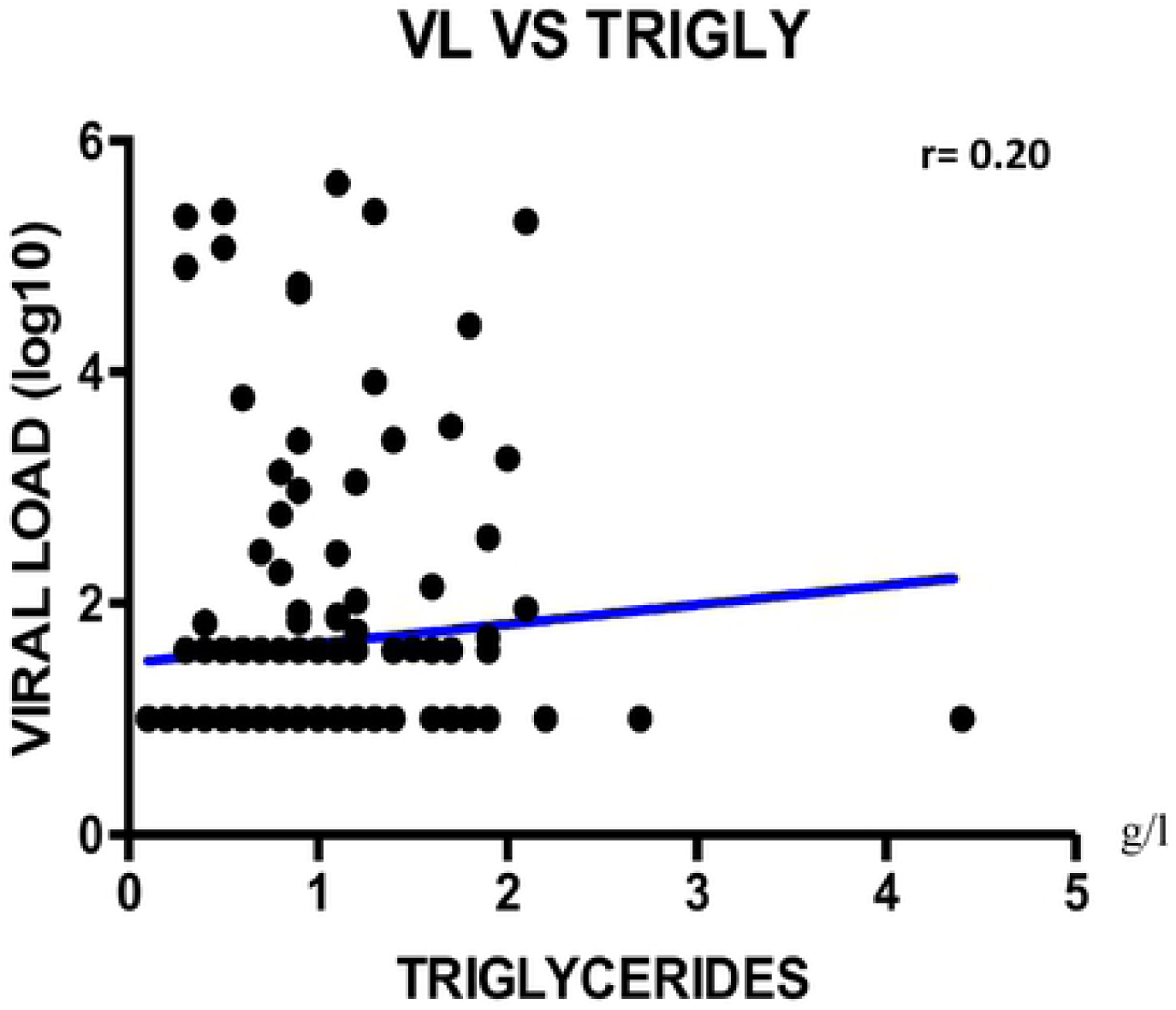
Correlation between viral load and total protein.

**Figure 6:**
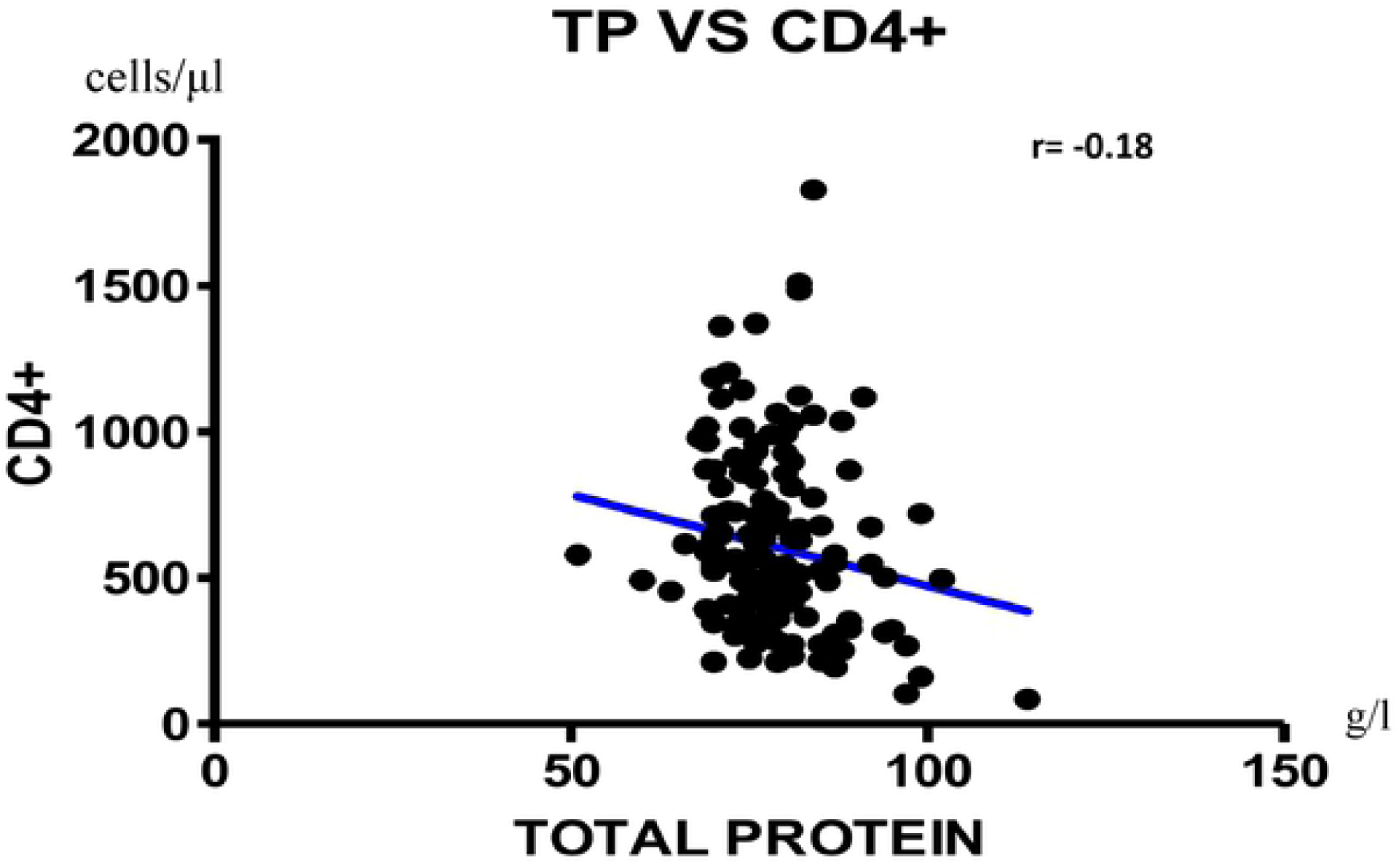
Correlation between viral load and triglycerides.

## Discussion

In the present study whose aim was to determine the correlation between the immuno-virological response and the nutritional profile of patients on ART within the Cameroonian context, the majority of our study participants were predominantly female (76.0%). This is similar to data in the same country reported by Ndawinz, *et al*. (68.1%), the Cameroon Population-based HIV Impact Assessment in 2017 (CAMPHIA) and the National Health and Demographic Survey (HDS) data in 2019 [17,18,27,28], thus confirming the biological vulnerability of women to HIV infection. The median age [IQR] of the study participants was 42 [33-51] years. This could be explained by the fact that the vast majority of infected individuals in Cameroon lie within this age interval, as found by Billong *et al*., in 2012 and previous survey data in Cameroon[17,18,20].

According to Body Mass Index (BMI), about 2 out of 5 participants were undernourished, suggesting that poor nutritional status is a major concern among patients receiving ART. Similar findings were recently reported in Ethiopia, which underscores the fact that beyond adherence to an effective ART-regimen, malnutrition remains a threat that deserves special considerations [29]. The fact that weight loss was found in 40% of our participants confirms the uniqueness of HIV infection in inducing a progressive decline of body weight [30,31]. Our data also revealed that about three quarters of participants lived on an average daily ration that range between $2 and $10 per day, and only half of participants could afford 2 meals per day, which in turn justifies the profile of WLP and highlights the financial constraint HIV could have on food insecurity among Cameroonian families [32].

About 79% of our participants had a normal level of total protein (66-88g/L), as previously reported in a similar African setting [29]. This profile also suggests upregulation of antibody production in an attempt to compensate ongoing immunodeficiency [33]. About half of our participants had a normal immunity (547 median CD4+ T cells), similar to previous reports in Cameroon [34,35]. The negatively weak correlation between CD4+ T cells and total protein was previously reported by Lyer *et al.* in a population of black Americans, with a similar positive trend observed with CD8 cells [36,37]. These states of immunity (CD4+ and CD8+) reflect the good response to ART [38]. The negatively weak correlation between CD8+ T cells and glucose suggests that HIV-associated alterations in CD8+ T-cell function also impair glycolysis, thus hindering glucose metabolism in HIV infection [39].

About 75% and 88% of participants achieved viral undetectability (<40%) and viral suppression (<1000 copies/mL) respectively. Similar to previous reports [34], these findings suggests that achieving the third pillar of the 90-90-90 is possible and could be reinforced with adequate nutrition in the frame of an effective ART [40].

Following multivariate analysis, total protein, total cholesterol and triglycerides were independent factors correlated to viral load. Of note, HIV infection could trigger a loss in total protein [41], which in turn lead to an increasing risk of proteinemia in the frame of an increasing viral load and CD4 count <200/mm^3^ [36]. Increased viral load is also known to impair cholesterol level [42], suggesting interaction between HIV and cholesterol leading to metabolic abnormalities (lipodystrophy, dyslipidemia, diabetes mellitus, and insulin resistance) prone by HIV itself and/or antiretroviral agents [43]. With a high correlation, total cholesterol thus appear to play an important role in HIV life cycle owing to lipid functions in viral entry, uncoating, replication, protein synthesis, assembly, budding and infectivity [44].

### Study limitations

Our study covered only a single region of Cameroon, making generalizability difficult at country level. It would have been insightful to assess pro and anti-inflammatory markers with regards to ART response and variability of the nutritional status. In addition, understanding the impact of immune activation induced by ART would have help in delineating covariates of nutritional parameters, immune response and accelerated ageing among patients receiving ART in our context.

## Conclusion

In this resources limited sitting, with patients having about five years of ART-experience, half are immunocompromised while the majority have achieved good virological response. Interestingly, one out of eight patients might be experiencing malnutrition. Specifically, increasing CD4 may favour hypo-proteinemia while increasing viral load may prone hyper-proteinemia and hypo-cholesterolemia. Further studies are needed in resources limited sitting with high burden of HIV-infection.

## Supporting Information

SI File (questionnaire)

## Tables and legends

**Table 1:** Correlation between immune-virological parameters and nutritional profile

|  | CD4 | CD8 | Viremia |
| --- | --- | --- | --- |
| Albumin | 0.50 | 0.93 | **0.013** |
| BMI | 0.80 | 0.78 | 0.05 |
| Calcium | 0.07 | 0.11 | 0.91 |
| Glucose | 0.87 | **0.003** | 0.38 |
| Iron | 0.10 | 0.65 | 0.68 |
| Magnesium | 0.31 | 0.34 | 0.76 |
| NRI | 0.45 | 0.66 | **0.0009** |
| Total Cholesterol | 0.89 | 0.08 | **0.0007** |
| Total protein | **0.03** | **0.031** | **0.0006** |
| Triglycerides | 0.58 | 0.46 | **0.018** |
| WLP | 0.46 | 0.89 | 0.19 |

**Legend:** BMI: Body Mass Index; NRI: Nutritional Risk Index; WLP: Weight Loss Percentage

**Table 2:**
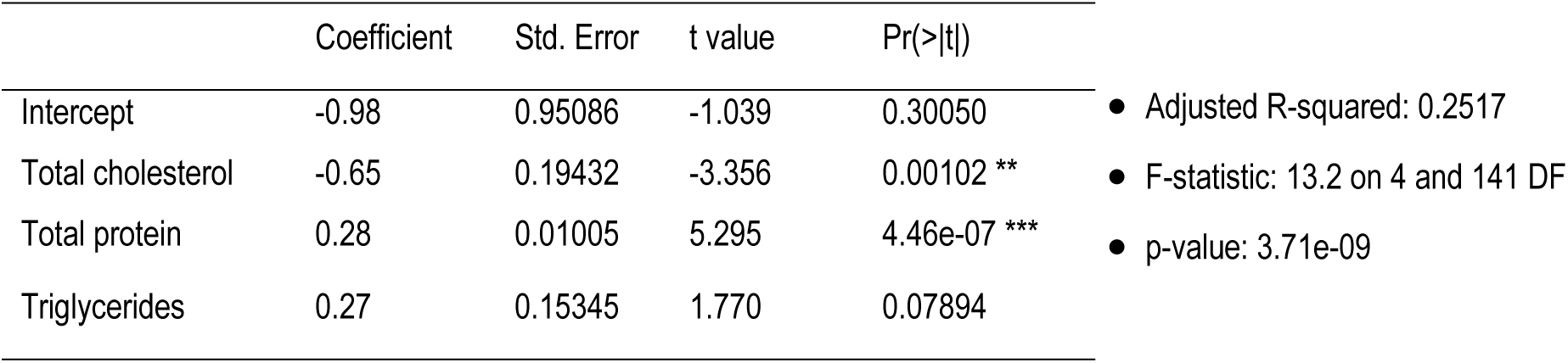
Multivariate analysis with viral load

## Acknowledgements

We are grateful to all HIV-clinic attendees at the care unit of the two different Health Facilities selected in the East region of Cameroon (Bertoua Regional Hospital and Nkolbikon Catholic Health Center). We are also thankful to the lab-technicians and clinicians of these sites who contributed in the field for enrolment; and the entire staff of the CIRCB for contributing in the lab-analysis process.

## Author Contributions

Designed the study: AA, RSK, JF, SMS, AN.

Collected the data: AA, ADN, ENS.

Analyzed and Interpreted the data: AA, ACK, MTT, ACC, JFY.

Initiated the manuscript: AA, RSK, JF, SMS, AN.

Revised the manuscript: ADN, ENS, ACK, MTT, ACC, YJ.

Approved the final version: All the authors.

